# Structural and Metabolic Insufficiency of the Blood-Brain Barrier Is Associated with Schizophrenia Penetrance

**DOI:** 10.64898/2026.11.18.767669

**Authors:** Frances M. Smith, Stewart A. Anderson, Jorge I. Alvarez

**Affiliations:** Department of Pathobiology, School of Veterinary Medicine, University of Pennsylvania, Philadelphia, Pennsylvania, United States; Department of Child and Adolescent Psychiatry, Children’s Hospital of Philadelphia, Philadelphia, Pennsylvania, United States; Department of Psychiatry, Perelman School of Medicine, University of Pennsylvania, Philadelphia, Pennsylvania, United States

**Keywords:** Blood-brain barrier, barrier integrity, neurovasculature, schizophrenia, 22q11.2 deletion syndrome

## Abstract

The blood-brain barrier (BBB) is a highly specialized vascular endothelial network tightly regulating peripheral influences on the brain. BBB endothelial cells (ECs) are mitochondrially rich compared to peripheral ECs, and mitochondrial function has been linked with the induction of BBB properties. In the 22q11.2 deletion syndrome (22qDS), a condition with heightened risk for schizophrenia (SZ), we demonstrated rescue of BBB properties by restoring mitochondrial function. Thus, we sought to understand if energetic and structural capacities of the BBB may influence the SZ penetrance seen in 22qDS. We differentiated induced pluripotent stem cells (iPSCs) from control, 22qDS individuals *without* SZ (22qDS(−)SZ), and 22qDS individuals *with* SZ (22qDS + SZ) into BBB-like cells (iBBB). We found that the iBBBs derived from 22qDS + SZ display considerable reductions in the rate of oxygen consumption as compared to 22qDS *without* (22qDS(−)SZ). Furthermore, 22qDS(−)SZ iBBB retain control-level barrier characteristics, including resistance to ionic flux, and enhanced tight junction and cytoskeletal properties. These phenotypes were further reinforced by RNA sequencing, which implicated several metabolic-, cytoskeletal-, and cell-cell adhesion-related differentially expressed genes, and ontological and gene-set enrichment phenotypes. Together, our functional and transcriptomic data indicate a dual compensation in metabolic and structural programs specific to the 22qDS subgroup lacking a SZ diagnosis. Thus, the penetrance of SZ in the BBB strongly associates with the capacity to maintain structural and metabolic phenotypes, representing a defining feature of 22qDS-related SZ pathophysiology. These findings bridge a gap in knowledge of molecular features which characterize differential phenotypes in the psychosis subtypes of 22qDS and address divergent BBB cell biology between individuals *with* and *without* SZ.

## 1. Introduction

Several developmental, intellectual, and psychiatric conditions are associated with aberrant mitochondrial function and blood-brain barrier (BBB) permeability [1–3], pathways which could represent a common theme underlying pathogenesis. Although neuronal dysfunction is recognized as a hallmark of such conditions, a neuro-centric perspective overlooks the neurovascular role in disease. Indeed, the BBB is a mitochondria-rich [4], cellular network that actively regulates the brain microenvironment by tightly regulating solute transport, waste clearance, and immune privilege [5–8]. Furthermore, mitochondrial function and barrier integrity are seemingly linked; exposing BBB monolayers to mitochondrial electron transport chain complex I and V inhibitors rapidly induces paracellular permeability and tight junction disorganization [9]. Thus, energetic weakness contributes to BBB permeability, and both are individually implicated in the disease pathogenesis of several central nervous system (CNS) diseases.

To investigate energetic and structural phenotypes of the BBB in CNS diseases, we leverage the human 22q11.2 deletion syndrome (22qDS), which confers haploinsufficiency of 46 protein-coding genes [10], including 7 related to mitochondrial physiology and function [11, 12], and the essential BBB tight junction protein Claudin-5 (Cldn5) [13]. 22qDS carries a heightened risk of intellectual disability and autism [14], and up to one quarter develop schizophrenia (SZ) [10]. This represents a 25-fold increase in SZ risk over the general population, making 22qDS an advantageous model to study disease penetrance. In induced pluripotent stem cells (iPSCs) differentiated to BBB-like (iBBB) cells from 22qDS + SZ individuals compared to healthy controls, we identified paracellular barrier permeability, disorganized junctional Cldn5 expression, compromised immune privilege [15, 16] and impaired oxidative phosphorylation [17]. Another group studying a different set of lines also found increased permeability and altered transcriptomics 22qDS + SZ iBBB relative to controls [18]. Additionally, we found that bezafibrate, a medication that enhances mitochondrial biogenesis, rescued BBB integrity in 22qDS + SZ iBBBs and 22qDS model mice (22qMc) [17]. In the mice, there was a robust negative correlation between the BBB leakiness and social memory, and bezafibrate also rescued this cognitive deficit despite being undetectable in the brains of 22qMc [17].

Since weaker mitochondrial function is associated with SZ penetrance in 22qDS iNeurons [19], enhancement of mitochondrial energetics reversibly influence BBB integrity and behavior [17], and CLDN5 haploinsufficiency is expected to influence BBB integrity in 22qDS [20], we hypothesize that the synergistic influences of structural and energetic compromise at the BBB influence SZ risk. To study this hypothesis, we studied iPSCs from healthy controls, 22qDS(−)SZ, and 22qDS + SZ individuals differentiated to iBBB cells [15, 17]. We found that in contrast to the iBBB from 22qDS + SZ, those from the 22qDS(−)SZ group have control levels of both oxidative phosphorylation and barrier function. In addition, RNA sequencing revealed that only the 22qDS(−)SZ group showed compensatory upregulation of pathways related to both mitochondrial energetics and cell-cell interactions, the mitochondrial effects being similar to compensation we previously described in 22qDS + SZ iNeurons [19]. These data uncover differential BBB physiology with respect to the SZ diagnosis in 22qDS despite their shared heterozygosity of 7 mitochondrial genes and Cldn5. We propose that structural and bioenergetic dysfunction of the BBB may contribute to SZ penetrance, positioning the neurovasculature as both a mechanistic driver of disease and a novel target for therapeutic intervention.

## 2. Materials and Methods

### 2.1 Cells

22qDS patient– and control-derived induced pluripotent stem cells (iPSCs), Table 1; kindly provided by Dr. Sergiu Pașca, Stanford University [21]) were differentiated to blood-brain barrier-like cells (iBBB) as in [22] and our previous studies [15, 17]. 22qDS cells were verified to have the deletion based on MLPA analysis [19] and construction of a heatmap containing the transcriptomic signature of all 46 genes in the A-D region (Supp. Fig. 1). Briefly, iPSCs were plated on Matrigel in complete StemMacs iPS-Brew supplemented with 10uM Y-27632 for 24-hours. For the following 6-days, media was replaced with “unconditioned media” (DMEM, 20% knock-out serum replacement, MEM non-essential amino acids, 200mM L-glutamine, 55mM Beta-mercaptoethanol, and Penicillin/Streptomycin cocktail). On day 7, media was replaced with “iBBB media” (HESFM + 0.5X B-27 without antioxidants, 10uM retinoic acid, and 20ng/ml bFGF). Additional iBBB media was added on day 8. On day 9, cells were lifted with Accutase and cryopreserved in 10% DMSO, 60% iBBB media, 30% FBS until the day of the assay.

**Figure 1.**
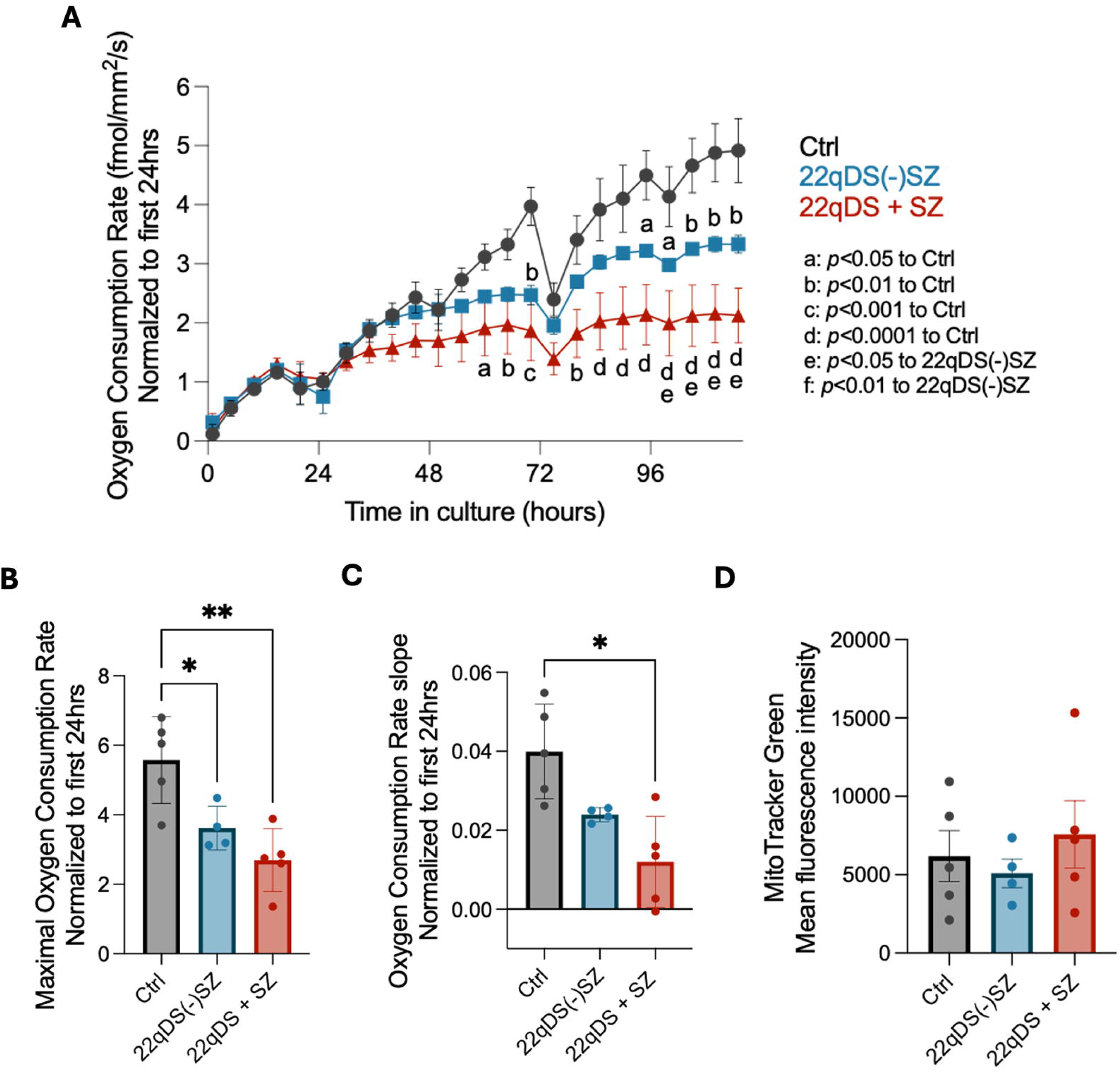
Mitochondrial metabolism in the SZ spectrum of 22qDS. (**A**) Oxygen consumption rate (OCR) was measured using the Lucid Labs RESHIPHER system for 120 hours in culture. Undulations in the trace result from media changes every 24-hours. (**B**) Maximal OCR for individual cell lines**. (C**) Slope of the linear regression of the individual lines**. (D**) Total mitochondrial content was assessed by flow cytometry using MitoTracker Green. *n* = 4-5 per cell lines per group with at least 2 technical replicates averaged C. (A) Two-way ANOVA, and (B-D) One-way ANOVA, each at α=0.05. *; *p*<0.05, **; *p*<0.01.

Cells were maintained at 37°C with 5% CO2 and 95% relative humidity throughout all experiments. Thawed iBBB were counted, plated on 40% human placental collagen IV, 10% bovine fibronectin, and cultured in iBBB media containing Y-27623. Cells were plated at 100,000 – 250,000 cells/cm^2^ for experiments described. 24-hours post-plating, media was replaced with iBBB media +B27 without other supplements. Media was replaced every 24-48 hours thereafter.

### 2.2 Oxygen Consumption Rate (OCR)

iBBB were plated in duplicate or triplicate in standard flat-well 96-well plates (Corning) and oxygen consumption was monitored for 120 hours using the Lucid Labs Resipher 32-well format oximetry system as previously shown [17]. Following the end of the experiment, wells were aspirated, washed once in PBS, lysed in protease inhibitor-containing RIPA buffer, and quantified for protein content using the BCA assay. Data from any sample falling statistically significantly below the average protein content of other wells in the experiment was discarded. Undulations in the oximetry readings reflect media changes. Data are represented as fold change from the first 24 hours in culture to study the maturation in aerobic metabolic function in an established monolayer. A linear regression model was fit to the traces of individual cell lines and the slope used to calculate metabolic maturation of the monolayer over time.

### 2.3 Mitochondrial quantification and flow cytometry

iBBB were plated at in 6-well plates and stained upon confluency. Cells were lifted using Accutase, washed once in DMEM, and resuspended in culture media containing 1μM MitoTracker Green. Cells were then incubated for 45 minutes at 37 degrees, shaking at 75 RPM. 0.2% Ghost Dye live/dead stain was spiked into the incubation, and the reaction allowed to continue shaking for 15 more minutes. Cells were spun at 300xg for 10 minutes and washed twice in PBS +1% FBS prior to running through the LSR Fortessa D cytometer (BD Biosciences). Mean fluorescence intensity of MitoTracker Green in live cells was quantified using FlowJo version 10 software.

### 2.4 Transendothelial electrical resistance (TEER)

iBBB were plated in duplicate or triplicate in 8W10E+ ECIS Z-Theta plates (Ibidi) as previously shown [17, 23]. Barrier integrity (resistance, 4k Hz) and substrate attachment (inversely correlated to capacitance, 64k Hz) were monitored for 120 hours. At the of the experiment, wells were aspirated, washed once in PBS, lysed in protease inhibitor-containing RIPA buffer, and protein content was quantified using the BCA assay. Data from any sample falling statistically significantly below the average protein content of other wells in the experiment was discarded. Data are represented as fold change from the first datapoint to analyze the maturation of barrier function. Area under the curve was calculated for the traces of the individual lines to derive the maintenance of the structural properties in the monolayer.

Correlation of the OCR and TEER are represented per cell line per group. A linear regression model was fit for each iBBB group, and the slope denoted. Pearson’s correlation was measured per group, and the *r* value represented. The BBB health index is the additive value of an individual cell line’s maximal respiration and maximal barrier integrity.

### 2.5 Immunofluorescence

iBBB were plated in 8-chamber glass slides (Invitrogen) and stained upon confluency as previously established [15]. In brief, cells were fixed with 3.7% paraformaldehyde for 10 minutes, washed twice in PBS, and permeabilized in 0.1% triton-X 100 for 10 minutes. Cells were washed twice in PBS and blocked in 10% normal donkey serum for 40 minutes, then incubated overnight with rabbit-anti-Cldn5 in 3% normal donkey serum. The following day, cells were washed thrice in PBS Tween (PBST) and incubated with donkey-anti-rabbit:488 and Phalloidin:647 for 40 minutes. Cells were washed thrice more in PBST, incubated with Hoechst nuclear stain for 10 minutes, washed thrice in PBST, and mounted in Mowiol mounting solution. Images were acquired using the Leica Thunder under 40X oil-immersion magnification.

#### Immunofluorescent quantification

Whole image pixel intensity of each stain (Cldn5 or phalloidin) was divided by the total pixel intensity in the Hoechst channel to obtain the average total staining intensity per cell. To measure staining presentation at cell junctions, regions of interest were drawn circumscribing the borders of 5-10 adjacent cells per image per cell line. The mean pixel intensity in that ROI was then divided by the *intensity per cell* quantified in the previous step, representing the fraction of total cell stain, which is present at the membrane, or the “junction-associated fraction.” 6-9 images were quantified per cell line. All analyses were conducted using ImageJ (NIH), while blinded to image identification.

### 2.6 RNA sequencing

iBBB were plated in 6-well plates, lysed upon confluency, and RNA was extracted according to the Qiagen RNAeasy Mini kit. Concentration and purity *via* A260/280 were verified using the ThermoFisher Nanodrop prior to sending 200ng RNA to Novogene for Illumina Sequencing. RNA integrity number of 9.8-9.9 for all samples was verified via bioanalyzer. To address the heterogeneity between individuals of the 22qDS group, we used DESeq2 with parameters set to *p-value*<0.01 and absolute value (Log2FoldChange)>1 to identify differentially expressed genes. From there, we ran these data against Hallmark, Reactome, KEGG Legacy and Medicus, Biocarta, and PID GSEA databases using 2,000 permutations under the *gene-set* program. Gene sets were considered significant under *p-value*<0.05 and FDR<0.05. Raw and processed data are uploaded to GEO under the accession number GSE353160.

### 2.7 Statistics

GraphPad Prism 10 was used for all statistical analyses, using α=0.05. Experiments were performed with at least 2 technical replicates. Data were averaged per cell line, and thus, the *n* value in each figure is representative of cell lines. Data are presented as mean +/-SEM.

## 3. RESULTS

### 3.1 Oxidative phosphorylation in iBBB cells distinguishes SZ diagnosis in 22qDS

We previously found that iBBB from 22qDS + SZ subjects are deficient in oxidative phosphorylation compared to healthy controls, and this is reversible by promoting mitochondrial biogenesis [17]. To determine whether this deficit is a property of 22qDS generally, or is particularly associated with 22qDS + SZ, we quantified the capacity for oxidative phosphorylation via oxygen consumption rate (OCR) over a 5-day period. Although after 36 hours the OCR tracked lower for both of the 22qDS groups relative to control, OCR of the +SZ group was consistently below that of the (−)SZ group (**Fig. 1A**). Analysis of OCR post 24 hours, normalized against the first 24 hours, revealed significant decreases relative to controls in both 22qDS groups (**Fig 1B**). However, the OCR trace for control and 22qDS(−)SZ iBBB climbs over time, indicating continually expanding metabolic activity of the established monolayer; versus the 22qDS + SZ iBBB whose metabolic activity remains virtually flat (**Fig. 1A).** This stagnation indicates impaired maturation of OCR in culture is specific to the 22qDS + SZ iBBB group (**Fig. 1C).**

Quantification of mitochondrial content revealed similar values among the three groups (**Fig. 1D**). Thus, while both 22qDS groups are deficient in OCR relative to controls, the difference is greater in the +SZ group especially at later time points, and this difference is not secondary to a reduction in mitochondrial mass.

### 3.2 BBB structural phenotypes diverge with respect to SZ penetrance in 22qDS

Since a 3’ UTR variant reducing Cldn5 levels on the remaining allele in 22qDS increases SZ risk, and BBB dysfunction is associated with idiopathic SZ [24, 25], we next sought to determine whether barrier dysfunction in 22qDS is selectively associated with SZ. We assessed transendothelial electrical resistance (TEER) to study barrier function and extracellular matrix (ECM) attachment, finding a deficit in paracellular barrier integrity exclusive to the 22qDS + SZ iBBBs (**Fig. 2**). On average, the 22qDS + SZ iBBB hit peak resistance earlier and at lower levels than either 22qDS(−)SZ or control iBBBs (whose traces were similar; **Fig. 2A**). While not statistically significant via one-way ANOVA, the 22qDS + SZ iBBB on average reach only 77% the maximal barrier function (**Fig. 2B**), and 72% barrier maintenance (quantified via area under the curve; **Fig. 2C**) of the 22qDS(−)SZ iBBB. Nevertheless, the 22qDS(−)SZ iBBB retain control-level barrier functions, indicating a strong association of barrier dysfunction in 22qDS + SZ to SZ comorbidity.

**Figure 2.**
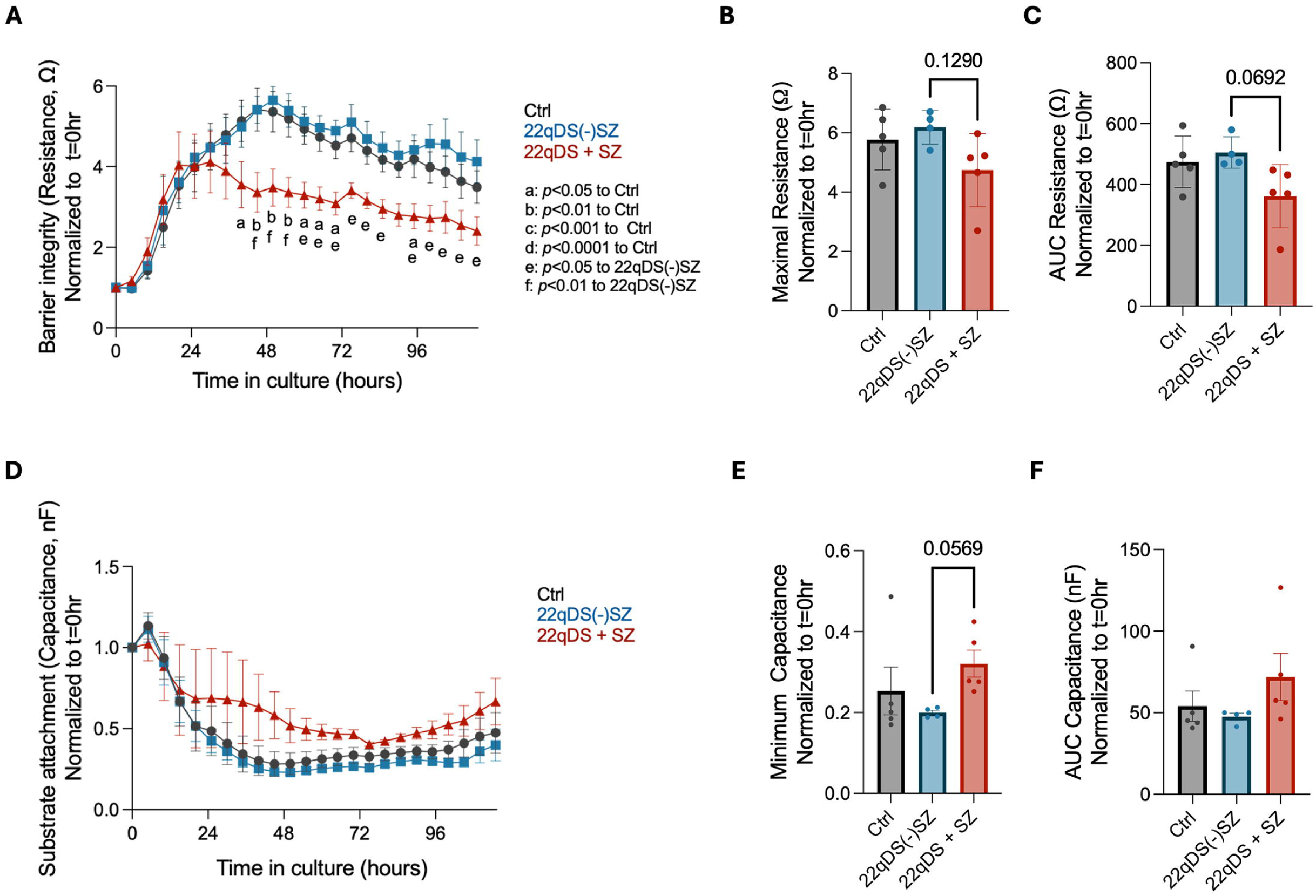
Barrier integrity and substrate attachment in iBBBs. (**A**) Barrier resistance (R, 4,000 Hz) was measured for 120 hours in culture. (**B**) Maximal barrier integrity reached by the individual cell lines. (**C**) Area under the curve for the individual lines. (**D**) Capacitance (C, 64,000 Hz) was measured using the ECIS Z-Theta TEER instrument for 120 hours in culture. (**D**) Minimal capacitance reached by the individual cell lines (**F**) Area under the curve for the individual lines. n = 4-5 cell lines per group with at least 2 technical replicates averaged. (A, D) Two-way ANOVA, and (B, C,E,F) One-way ANOVA, each at α=0.05.

We then examined the ability of these cells to maintain ECM attachment, for which capacitance values are inversely correlated (**Fig. 2D).** In contrast to barrier resistance (**Fig. 2A**), the substrate attachment properties do not reflect a meaningful time-dependent difference between the three iBBB groups. While the minimum capacitance reached between the 22qDS subgroups approaches statistical significance, the ability to maintain ECM attachment is unaffected (**Fig. 2E**). Taken together, the 22qDS(−)SZ iBBB retain control-level barrier integrity and ECM attachment capacities, while the 22qDS + SZ iBBBs only retain ECM attachment but not paracellular integrity. These results reveal differential barrier physiology in 22qDS dependent upon SZ penetrance.

### 3.3 Cldn5 expression and organization differ according to SZ penetrance in 22qDS BBB

In addition to the inclusion of Cldn5 in the 22qDS deleted region, a 3’ UTR Cldn5 variant results in reduced protein expression [20], and is associated with SZ risk in females [25]. These findings further implicate Cldn5 in SZ pathogenesis, and thus we sought to assess the expression and organization of Cldn5 in our iBBBs with respect to the SZ diagnosis. Cldn5 shows aberrant staining patterns at multi-cellular contacts in the 22qDS + SZ iBBBs (**Fig. 3A**), representing frayed and disorganized patterning. We have observed this theme in the past when comparing 22qDS + SZ iBBB relative to healthy controls [15], and it contrasts to the cobblestone-like junctional staining pattern characteristic of iBBBs [22], which is observed in both the control and 22qDS(−)SZ groups. This may reflect vulnerability of the plasma membrane’s physical and mechanical properties in the 22qDS + SZ iBBB. As Cldn5 is only functional at adjacent EC interfaces, we sought to quantify how much of this protein was presenting at cell-cell borders. Interestingly, this revealed that the 22qDS(−)SZ iBBB have increased junction-associated Cldn5 compared to both the control and 22qDS + SZ iBBBs (**Fig. 3B**). This is reinforced by heightened cytoplasmic staining in the 22qDS + SZ iBBB (**Fig. 3A**), and may represent Cldn5 internalization. In total, the 22qDS(−)SZ iBBBs express a higher fraction of Cldn5 at cell junctions, and retain normal Cldn5 staining morphology, both of which could contribute to the differences observed in barrier function between this group and 22qDS + SZ. (**Fig. 2A-C).**

**Figure 3.**
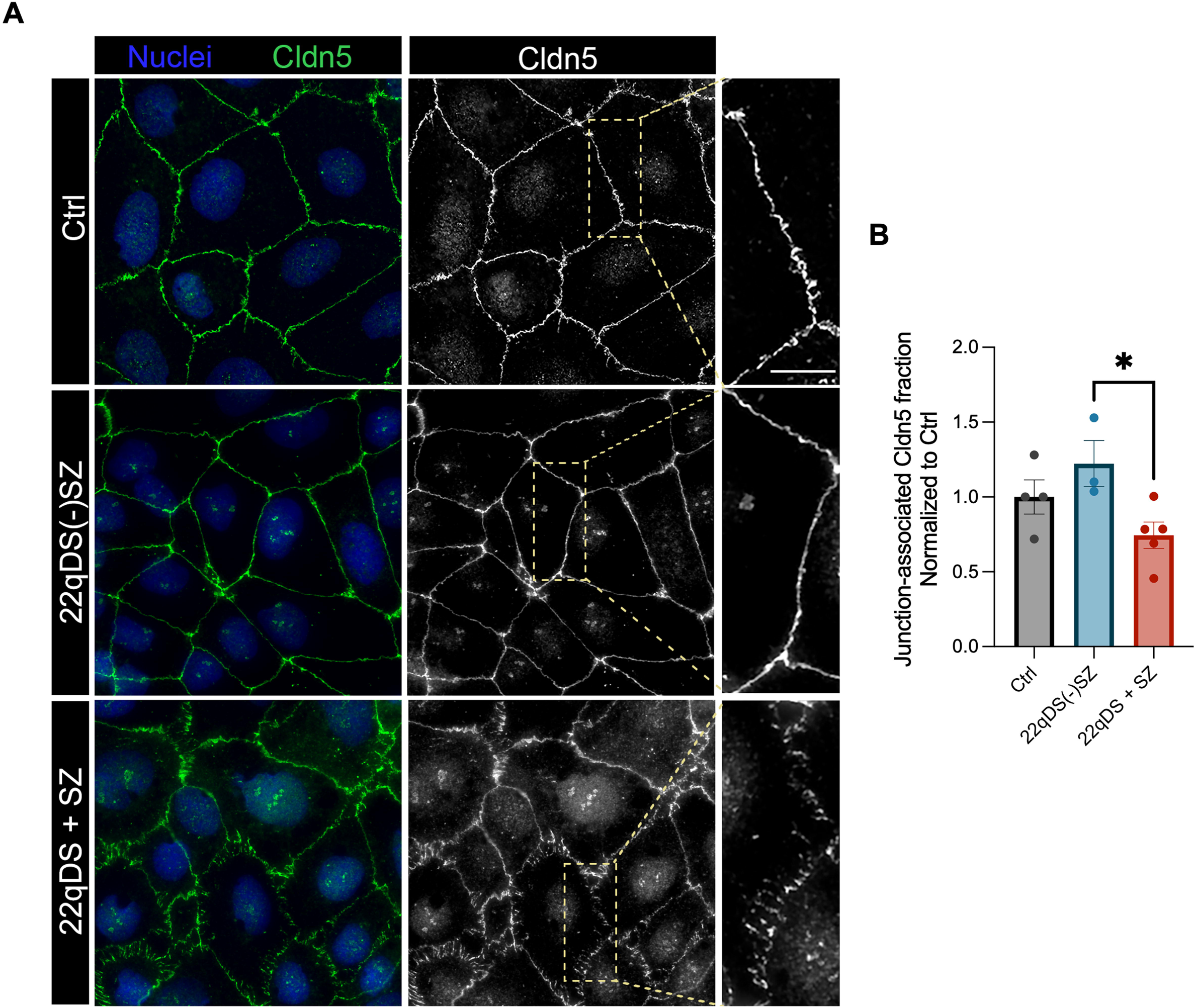
Cell-cell junction Cldn5 expression. (**A**) Representative iBBB monolayers co-stained with Cldn5 (green) and Hoechst nuclear dye (blue). Scale bar, 20μm. (**B**) Junction-associated Cldn5 fraction. n = 3-4 cell lines per group. One-way ANOVA at α=0.05. *; p<0.05.

### 3.4 F-actin junctional delineation differentially associates with SZ in 22qDS BBB

Actin is the most abundant cytoskeletal protein, and its anchorage of tight junction proteins including Cldn5 is essential to maintaining barrier function [26]. Thus, we sought to examine total, and membrane-associated filamentous actin (F-actin) abundance in our iBBBs (**Fig. 4A**). Consistent with impaired barrier integrity (**Fig. 2**) and decreased Cldn5 membrane association (**Fig. 3**) in 22qDS + SZ iBBB, we also expected a reduction in the association of F-actin with the membrane. Indeed, this is reinforced by the more well-defined plasma membrane delineation in the control and 22qDS(−)SZ compared to the 22qDS + SZ (**Fig. 4A**). Again, the 22qDS(−)SZ iBBBs have a stronger fraction of the total stain organized at cell junction interfaces compared to 22qDS + SZ (**Fig. 4B).** Thus, iBBB 22qDS + SZ are distinguished from the control and 22qDS(−)SZ by the impaired capacity to organize F-actin at adjacent cell borders. Collectively, combined reductions in junctional Cldn5 presentation (**Fig. 3),** and the underlying scaffolding supportive of tight junction function (**Fig. 4**) between adjacent cells of the iBBB monolayer likely underscores the increased barrier vulnerability (**Fig. 2**) of 22qDS + SZ iBBB.

**Figure 4.**
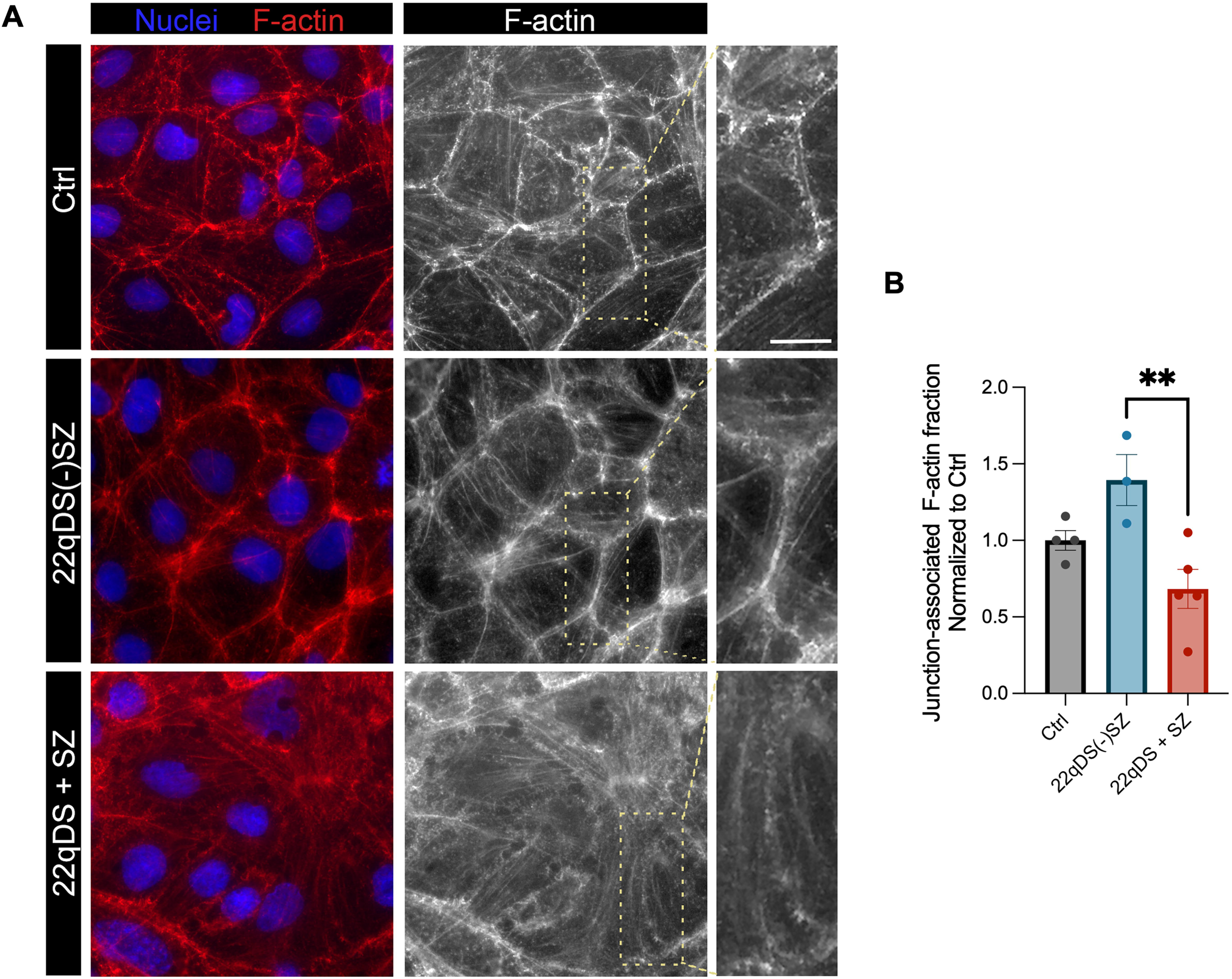
Association of F-actin at the cell-cell junction. (**A**) Representative iBBB monolayers co-stained with Hoechst nuclear dye (blue) and Phalloidin (red) to stain F-actin. Scale bar, 20μm. (**B**) Junction-associated F-actin fraction. n = 3-4 cell lines per group. One-way ANOVA at α=0.05. **; p<0.01.

### 3.5 SZ diagnosis in the 22qDS BBB is associated with altered mitochondrial and structural gene expression

Based on the striking differences in energy metabolism (**Fig. 1**) and cell structure (**Figs. 2-4**) defined by the SZ diagnosis in 22qDS, we performed RNA sequencing to examine the differential transcriptomic signatures associated with each group. Among the 3 iBBB groups (**Fig. 5A),** we identified 903 differentially expressed genes between control and 22qDS(−)SZ, 613 between control and 22qDS + SZ, and 221 between 22qDS(−)SZ and 22qDS + SZ (**Fig. 5B**). Based on the functional metabolic (**Fig. 1**) and structural (**Figs. 2-4**) similarity between control and 22qDS(−)SZ iBBB, we analyzed gene co-expression between these two groups, and found 90 shared differentially expressed genes enriched over the 22qDS + SZ iBBB (**Fig. 5C**). Of these 90 genes, several have pro-mitochondrial or structural-related functions. For instance, Mitoregulin (*MTLN*), an emerging gene at the nexus of several mitochondrial homeostasis pathways [27], and Adducin-2 (*ADD2*) an actin-binding protein critical to barrier integrity, and tight– and adherens-junction continuity [28].

**Figure 5.**
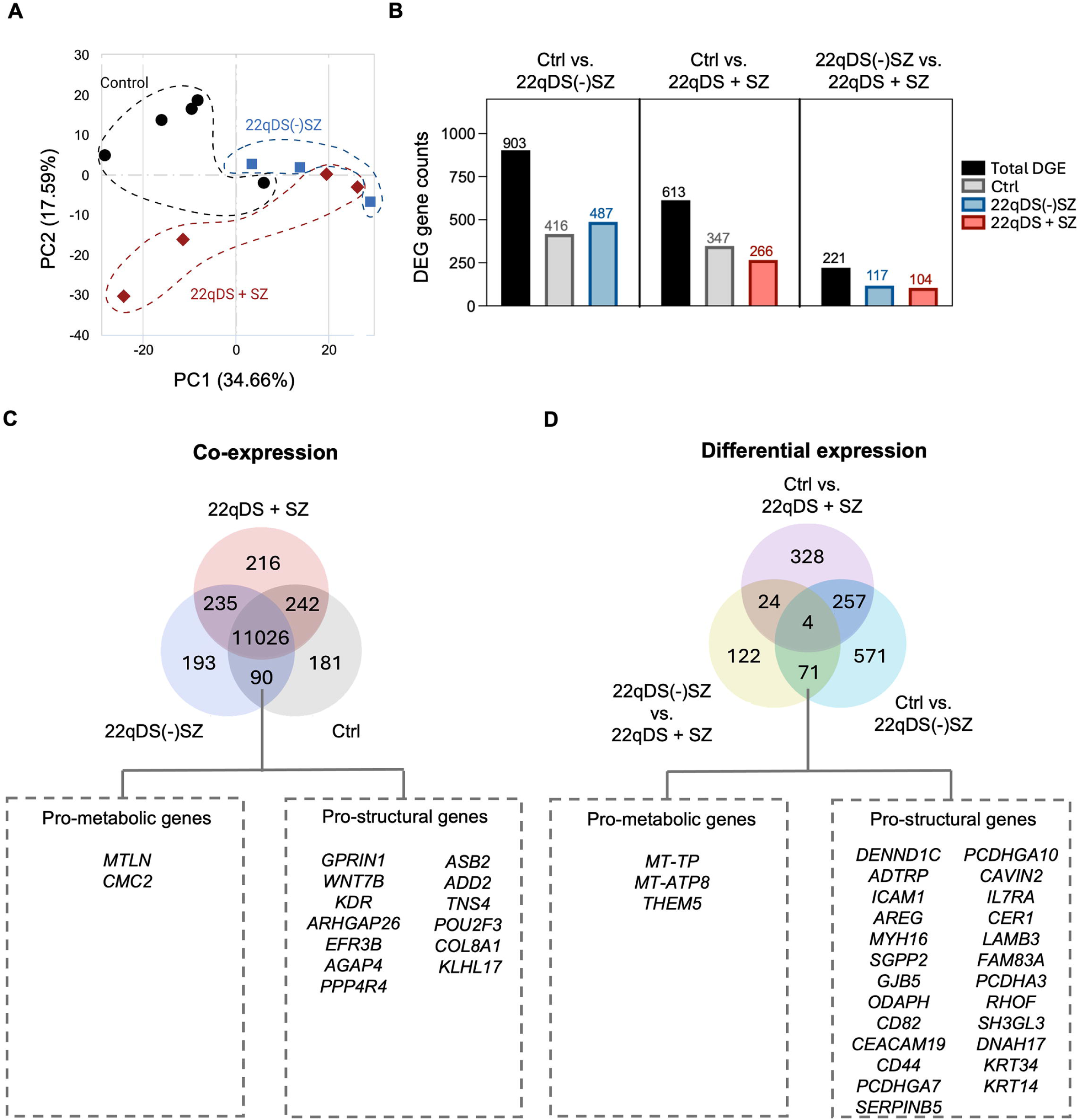
Gene expression in the iBBB groups. (**A**) PCA analysis of the three iBBB groups. (**B**) Numbers of differentially expressed genes (DEGs). (**C**) Gene co-expression; relevant mitochondrial and structural genes among the 90 DEGs expressed in Ctrl and 22qDS(−)SZ are annotated. (**D**) DEGs shared among the three comparison groups. Relevant mitochondrial and structural genes among the 71 DEGs upregulated in 22qDS(−)SZ relative to *both* Ctrls and 22qDS + SZ are annotated. *n* = 3-5 cell lines per group.

The differential gene expression enriched only in the 22qDS(−)SZ compared to both control and 22qDS + SZ further suggests a strong metabolic and structural compensatory phenotype exclusive to this group. This resulted in 71 genes enriched in the 22qDS(−)SZ iBBB (**Fig. 5D**), again including several mitochondrial and structural genes. Among them, two mitochondrially-encoded transcripts: *MT-TP,* a mitochondrial tRNA, and *MT-ATP8*, a subunit of the electron transport chain complex V, and *THEM5*, which localizes to the mitochondria for fatty acid metabolism. Many structural genes upregulated in 22qDS(−)SZ are pro-vascular, including *ADTRP* [29] and *AREG* [30], and several are structurally-supportive, such as the actin dynamics-accessory proteins *DENND1C* [31] and *RHOF* [32]. Therefore, the transcriptomic signature of 22qDS(−)SZ iBBB implicates structural and metabolic compensation, and conversely, vulnerability in 22qDS and SZ comorbidity. Therefore, these two biological axes, and their interactions may be valuable targets for diagnostic or therapeutic potential.

### 3.6 Divergent metabolic and structural transcriptomic signatures dependent upon SZ penetrance in 22qDS BBB

To understand the significance of gene-gene interactions and pathway enrichment in our samples, we ran gene ontology (GO) analysis (**Fig. 6A, D**). Intriguingly, within the top 5 BP pathways enriched in 22qDS(−)SZ iBBB, several include features of cell-cell adhesion, when compared to both 22qDS + SZ iBBB (**Fig. 6A**) and control iBBB (Supp. Fig. 2A), as well as plasma membrane-associated annotations (i.e., Clathrin-coating). Furthermore, Hallmark gene-set enrichment analyses (GSEA) annotations include strong enrichment in “oxidative phosphorylation” and “apical junction” pathways between 22qDS(−)SZ and 22qDS + SZ iBBB (**Fig. 6B**). Additional analyses using the Reactome, Kegg Medicus and Legacy, Biocarta, and Protein Interaction Database (PID) databases further revealed a robust number of differentially regulated pathways. As validation for our biochemical experiments, we emphasize those which are related to energetic and structural programs (**Fig. 6 C, F**), but a complete list of ranked GSEA pathways above a normalized enrichment score (NES) of 1.5 is represented in (Supp. Data file 1). These analyses revealed a strong enrichment in mitochondrial-protective pathways in the 22qDS(−)SZ relative to 22qDS + SZ, including mitophagy and mitochondrial biogenesis, mitochondrial translation, and oxidative phosphorylation. Additionally, structural pathways were emphasized, including programs for both junctional (tight, adherens, and gap junction) integrity as well as F-actin dynamics (with “Rho GTPase cycle” being the *second* most enriched pathway, behind “translation”) (Fig. 6C, Supp. Data file 1).

**Figure 6.**
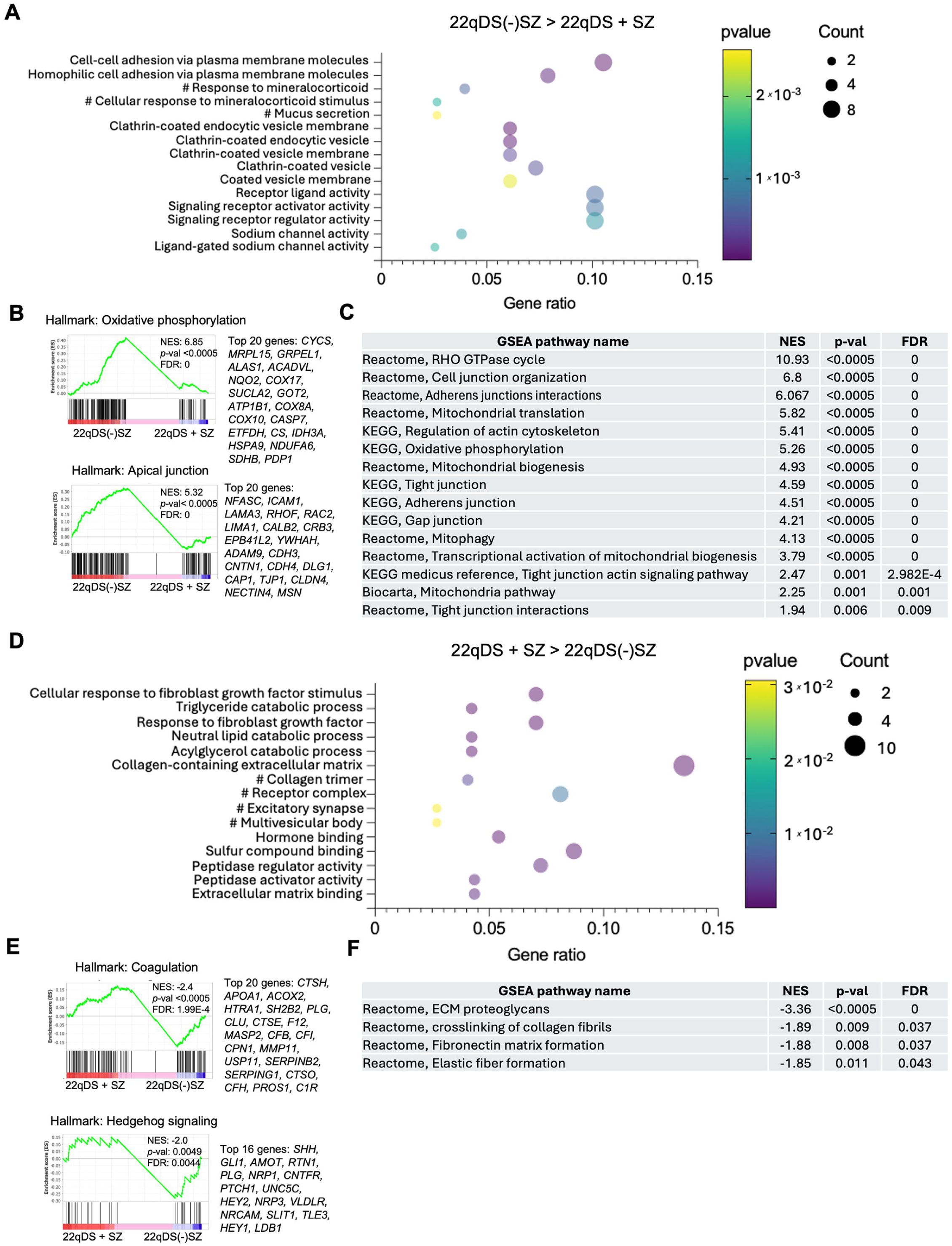
Metabolic and structural transcriptomic programs in the iBBBs. *Enrichment in 22qDS(−)SZ over 22qDS + SZ for* (**A) GO terms;** The top 5 significant gene ontology (GO) terms from biological process, cell compartment, and molecular function, respectively. (**B) Hallmark GSEA;** Select significant Hallmark GSEA annotations represented with the top 20 core enriched genes included to the right. (**C) Curated GSEA;** Select significant GSEA annotations from Reactome, KEGG or Biocarta databases. *Enrichment in 22qDS + SZ over 22qDS(−)SZ for* (**D) GO terms, (E) Hallmark,** and (**F) Curated GSEA.** #; GO terms which did not also fall below *p-adjusted value* of 0.1. *n* = 3-5 cell lines per group,

In contrast, the 22qDS + SZ iBBB have upregulated GO terms associated with cell*-substrate* interactions, particularly collagen-associated interactions, compared to both 22qDS(−)SZ (**Fig. 6D**) and control iBBB (Supp. Fig. 2B). This suggests differential compensatory pathways among the 22qDS subgroups, and reinforces the deficit in barrier resistance unique to 22qDS + SZ iBBB (**Fig. 2A-C**), but retention of capacitance (**Fig. 2D-F).** Hallmark GSEA pathways returned fewer programs but included gene sets associated with “coagulation” and “hedgehog signaling,” perhaps as products of an “activated” endothelial phenotype following an inflammatory or stress-induced state [15, 16] (**Fig. 6E**). Furthermore, the additional GSEA databases returned fewer structural pathways, and no mitochondrial programs enriched in 22qDS + SZ iBBB over 22qDS(−)SZ iBBB (**Fig. 6F).** The structural pathway hits detail cell-substrate interactions (ECM proteoglycans, collagen crosslinking, and fibronectin formation), again reinforcing differential structural phenotypes between the psychoses subtypes of the 22qDS iBBB, whereby the 22qDS(−)SZ strongly conserve cell-cell interactive pathways, versus the 22qDS + SZ which maintain cell-substrate interactions (**Figs. 2 and 6**). Unsurprisingly for a disease of chromosomal deletion, the top GO terms and GSEA annotations enriched in the control iBBB compared to both 22qDS subtypes include pathways associated with chromosomal division (Supp. Fig. 2 C, D). In total, the transcriptome drastically varies with respect to the SZ penetrance of 22qDS, pointing to the BBB’s capacity for energetic and structural maintenance as factors associated to psychiatric pathogenesis.

### 3.7 Contrasting structural-energetic correlations define the 22qDS BBB with respect to the SZ diagnosis

We previously demonstrated that enhancement of mitochondrial biogenesis promotes several features of barrier function in 22q mice and 22qDS + SZ iBBB compared to healthy controls, including barrier integrity, Cldn5 expression, and Cldn5 membrane organization [17]. Thus, given the differential metabolic and structural functions uncovered by our functional and transcriptomic analyses, we sought to demonstrate a correlative relationship between these two essential axes of BBB biology. Correlation of maximal OCR (**Fig. 1B**) and barrier resistance (**Fig. 2B**) per cell line identified a strongly positive linear correlation in the 22qDS(−)SZ group, and by contrast, a robustly negative linear correlation in 22qDS + SZ iBBBs (**Fig. 7A**). These data were used to derive the “BBB health index,” which demonstrates insufficiency exclusive to the 22qDS + SZ iBBB (**Fig. 7B**). Together, this suggests dual metabolic and structural compromise exclusive to the 22qDS + SZ group, and a unique synergistic program leveraged by the 22qDS(−)SZ group.

**Figure 7.**
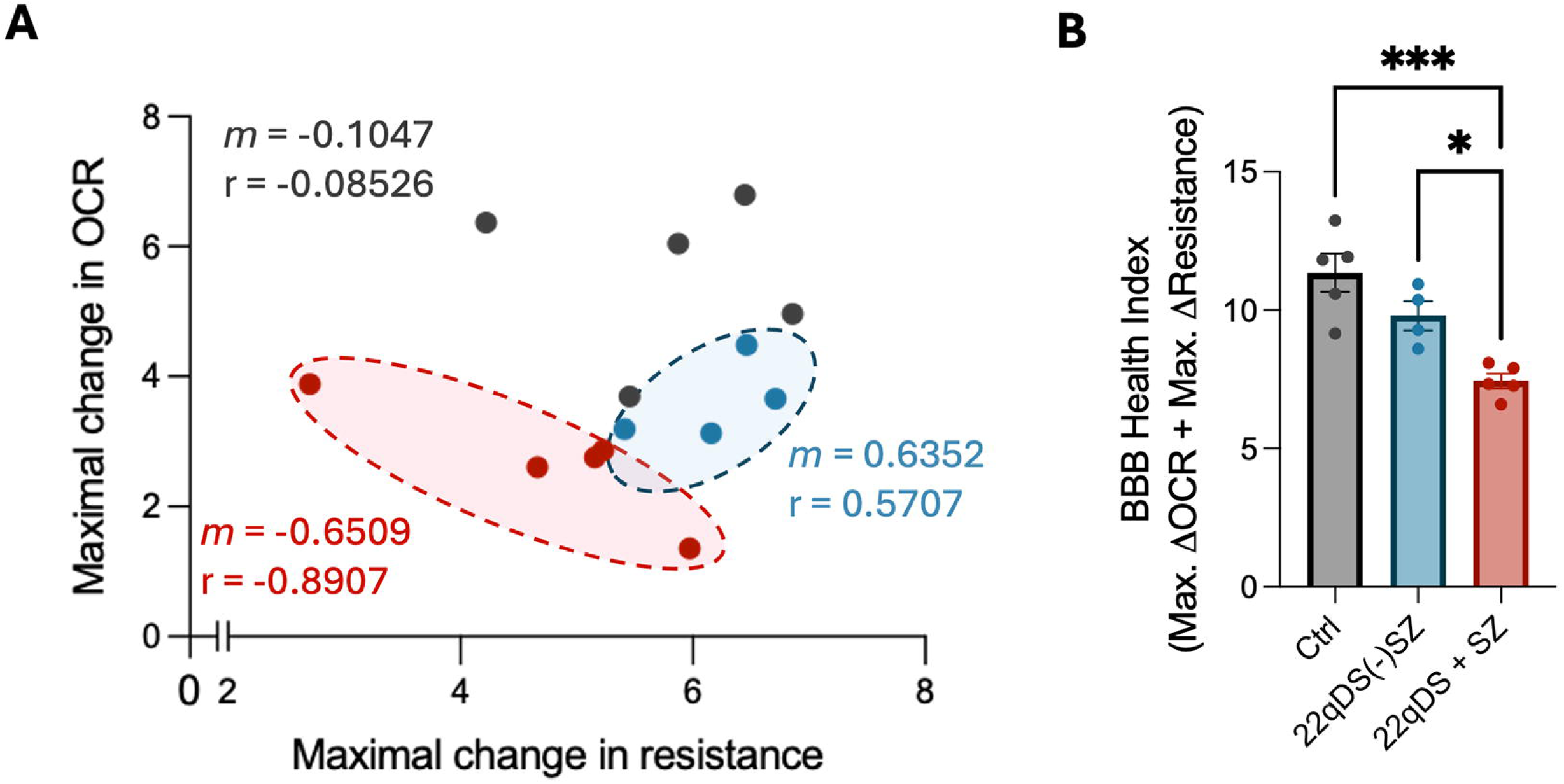
Correlation of structural and metabolic phenotypes across iBBB. (**A**) The maximal change in OCR (Fig. 1B) is plotted against the maximal change in barrier resistance (Fig. 2B) for individual cell lines. Slope (m) of the linear regression and Pearson’s correlation (r) are represented per iBBB group. (**B**) The “BBB health index;” the additive value of maximal OCR and barrier functions represent. n = 4-5 cell lines per group. (B) One-way ANOVA at α=0.05. *; p<0.05, ***; p<0.001.

Collectively, our data suggest that 22qDS(−)SZ iBBB largely retain control-level physiology in energetic and structural aspects; mild reduction in oxidative phosphorylation (**Fig. 1**), retention of paracellular barrier function (**Fig. 2**), and junctional integrity (**Figs. 3 and 4**), and upregulated energetic and structural transcriptomic programs (**Figs. 5 and 6**). On the contrary, 22qDS + SZ iBBB are deficient in *all* of these phenotypes.

## 4. DISCUSSION

The role of the BBB in the pathogenesis of CNS diseases is increasingly being appreciated, yet less is known about how the energetics at this critical interface support barrier function. Mitochondrial dysfunction and BBB permeability are individually implicated risk factors for diverse CNS pathologies, including psychoses [33, 34], which affects up to 25% of the 22qDS population. Previously, we identified that mitochondrial energetic deficits associates with SZ penetrance in 22qDS neurons and lymphoblastoid cell lines [19]. Motivated by the link between mitochondrial function and barrier integrity [9, 17], here we examined whether such deficits occur in the BBB. Our findings identify both structural and metabolic dysfunction as a hallmark of BBB pathology in 22qDS-associated SZ, expanding the framework of disease pathogenesis beyond neural deficits to include neurovascular mechanisms.

Since BBB dysfunction in 22qDS is mitigated by promoting mitochondrial biogenesis [17], and SZ differentially associates with the capacity for oxidative phosphorylation, we evaluated barrier integrity with respect to the SZ diagnosis. We found that the 22qDS(−)SZ iBBB display control-level barrier integrity while SZ comorbidity strongly associates with barrier dysfunction. Preservation of cell-cell interactions may be a defining feature of the 22qDS(−)SZ iBBB as this group upregulates Cldn5 and F-actin at adjacent cell contacts, and transcriptionally returns a strong theme for cell-cell interactions and junctional maintenance. Thus, 22qDS(−)SZ iBBB maintain several features related to structural function and barrier integrity. Given the clinical association of SZ and BBB permeability [2, 35], paracellular BBB breakdown may be an active influencer of SZ pathogenesis. Notably, cell adhesion defects in 22qDS iPSC-derived astrocytes parallel with mitochondrial dysfunction [36], supporting our findings of convergent structural and metabolic abnormalities driving BBB dysfunction in 22qDS and lending a potentially valuable therapeutic approach.

Mitochondrial function in BBB physiology is a relatively new area of study, but mitochondrial activity supports BBB properties [9, 17]. Hence, the retention of mitochondrial function in 22qDS(−)SZ iBBB likely supports barrier integrity. Indeed, we find that the 22qDS(−)SZ iBBB display a positive correlation between the metabolic and structural phenotypes measured here, which is not true of the 22qDS + SZ iBBB group. Notably, the 22qDS(−)SZ do still have a moderate reduction in the capacity for oxidative phosphorylation relative to control iBBB, while still mounting proper barrier integrity.

One explanation for this dichotomy is response to reactive oxygen species (ROS). Oxygen radicals are natural byproducts of oxidative phosphorylation [37], elevated in 22qDS models in part due to haploinsufficiency of the mitochondrial ROS scavenger thioredoxin reductase (TXNRD2) [38], and redox stress is a known contributor to BBB dysfunction [39, 40]. SZ itself is increasingly associated with ROS imbalance [41, 42], for instance, there are decreased levels of the antioxidants free glutathione and superoxide dismutase (SOD) in SZ patient blood [43], and both are protective of BBB function [44, 45]. Thus, the additional redox stress imposed by comorbid SZ in 22qDS may potentiate barrier dysfunction. An additional possibility is that more productive aerobic energy metabolism in the 22qDS(−)SZ iBBB provides the ATP necessary for f-actin metabolism. This concept is supported by the strong abundance of F-actin in the 22qDS(−)SZ iBBB, and the upregulation of genes regulating cytoskeletal dynamics in this group. Deeper investigation of this notion is a future direction of this work.

We report that 22qDS(−)SZ iBBB display several features of energetic resilience as compared to the 22qDS + SZ, similar to our previous report in iNeurons [19]. Linking our previous investigations detailing metabolic compromise of 22qDS + SZ compared to control iBBB [17], these data implicate that mitochondrial energy metabolism more strongly associates with the SZ diagnosis than the 22qDS deletion. This could suggest that SZ risk is associated with increased energetic vulnerability in 22qDS, and that 22qDS(−)SZ iBBB, like the iNeurons [17], leverage compensatory pro-mitochondrial programs. Indeed, our transcriptomic analyses support these contentions as the 22qDS(−)SZ iBBB upregulate transcriptional programs involved in mitochondrial biogenesis and mitophagy. Taking into consideration that we find similar mitochondrial content among the iBBB groups, this points to enhanced mitochondrial turnover in 22qDS(−)SZ as a mechanism to maintain a younger and more functional mitochondrial pool.

Our study has several limitations. First, our relatively low number of cell lines per group and the degree of clinical heterogeneity present in both 22qDS and SZ. Second, we report that energetic and structural compromise appear to *associate* with the SZ diagnosis, but we cannot distinguish cause from effect. As enhancing mitochondrial biogenesis rescues barrier dysfunction in 22qDS + SZ iBBBs and social memory deficits in 22qDS mice [17], these phenotypes appear to be reversible by enhancing mitochondrial function. Third, although 22qDS + SZ iBBBs show reduced membrane localization of Cldn5, and bezafibrate restores both Cldn5 localization to the plasma membrane and barrier function in 22qDS + SZ iBBB [17], it remains unclear whether the observed BBB defects arise primarily from impaired energetics, Cldn5 allelic insufficiency, or both. However, the finding that Cldn5 haploinsufficiency in mice affects BBB function [20], coupled with the increased SZ risk conferred by a variant in the 3’UTR on the remaining Cldn5 allele [20, 25], strongly suggests that both barrier structural weakness and impaired energetics contribute to the 22qDS + SZ BBB phenotype.

## 5. CONCLUSION

We sought to characterize differences among 22qDS iBBB with respect to the SZ diagnosis, finding that the 22qDS(−)SZ iBBB *functionally* mirror the energetic and structural functions of control iBBB, and *transcriptionally* compensate with dual energetic and structural programs while the 22qDS + SZ iBBB remain largely deficit in these respects. Thus, the SZ diagnosis is associated with robust energetic and structural compromise in iBBBs, consistent with energetic and neurovascular vulnerability in SZ [2, 34]. These findings support a model by which neurovascular bioenergetic and structural dysfunction contributes to SZ risk.

## FUNDING

This work was supported by the National Institute of Mental Health R01MH134893-01 to JIA and SAA, R01MH110185-03 to SAA, R01MH066912 to SAA, and R01MH134797-01 to JIA, the National Institute of Health grant to the Institute of Translational Medicine and Therapeutics (ITMAT) and by the Transdisciplinary Awards Program in Translational Medicine and Therapeutics (TAPITMAT) of the University of Pennsylvania NCATS 5UL1TR001871-04 to JIA and SAA, the Penrose Family to SAA, and the Uytengsu-Hamilton 22q11 Neuropsychiatry Research Program of the Maternal and Child Health Research Institute at Stanford University UH22QEXTFY22-03 to JIA.

## Acknowledgements

We acknowledge Katherine E. Benson (Summer undergraduate researcher) for her participation in the early stages of this project. We thank Dr. Amit Bar-Or for use of the Fortessa D cytometer along with the Penn Cytomics Core which maintains cytometer care. We thank the PennVet Imaging Core and the core manager, Dr. Ruthel Gordon for training and support in immunofluorescent detection, as well as Dr. Ethan Goldberg for access to the Lecia Thunder for which immunofluorescent images were acquired. Finally, we are indebted to Dr. Mark Khan for use of the ECIS TEER instrument.

## Abbreviations

22qDS: 22q11.2 deletion syndrome
22qMc: 22q mice
22qDS + SZ: 22qDS individuals with schizophrenia
22qDS(−)SZ: 22qDS individuals without schizophrenia
ADD2: Adducin-2
BBB: blood-brain barrier
CNS: central nervous system
EC: endothelial cell
ECM: extracellular matrix
F-actin: filamentous actin
GO: gene ontology
GSEA: gene-set enrichment analysis
iBBB: iPSC-derived BBB cell
iPSC: induced pluripotent stem cell
MTLN: Mitoregulin
NES: normalized enrichment score
OCR: oxygen consumption rate
PID: protein interaction database
ROS: reactive oxygen species
SOD: superoxide dismutase
SZ: schizophrenia
TEER: transendothelial electrical resistance
TXNRD2: thioredoxin reductase

## FIGURE LEGENDS

**Supplemental Figure 1.**
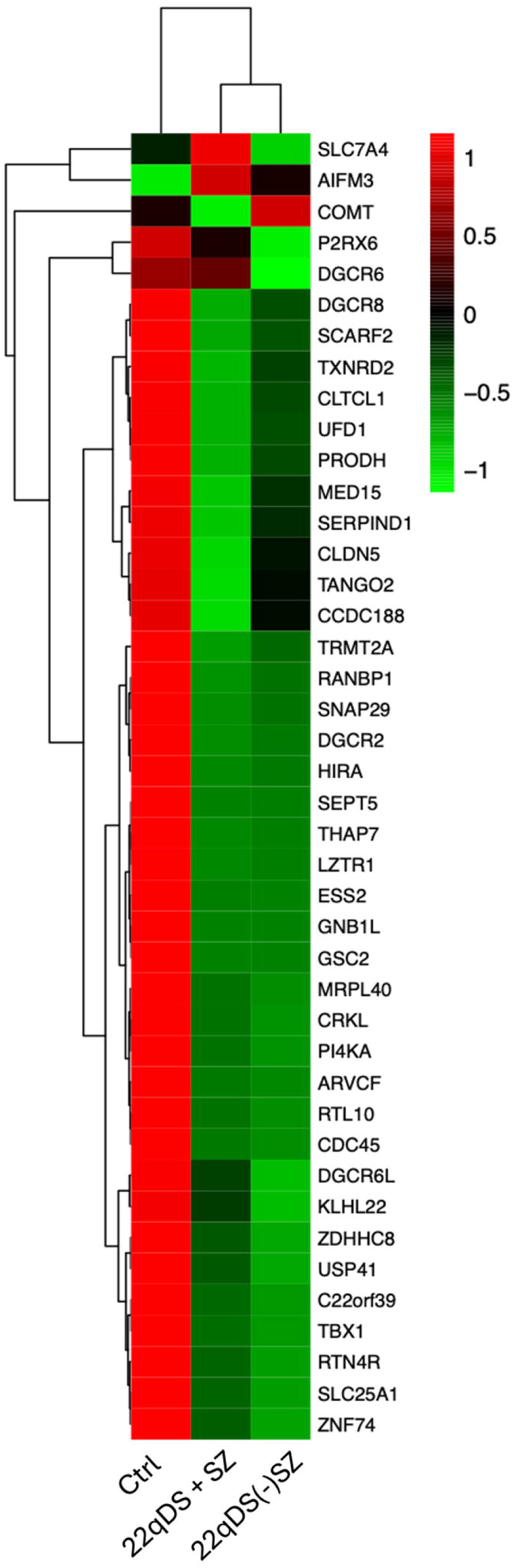
Verification of 22qDS in our iBBBs. Heatmap containing the 46 genes included in the A-D region of 22qDS. Red; high expression, green; low expression.

**Supplemental Figure 2.**
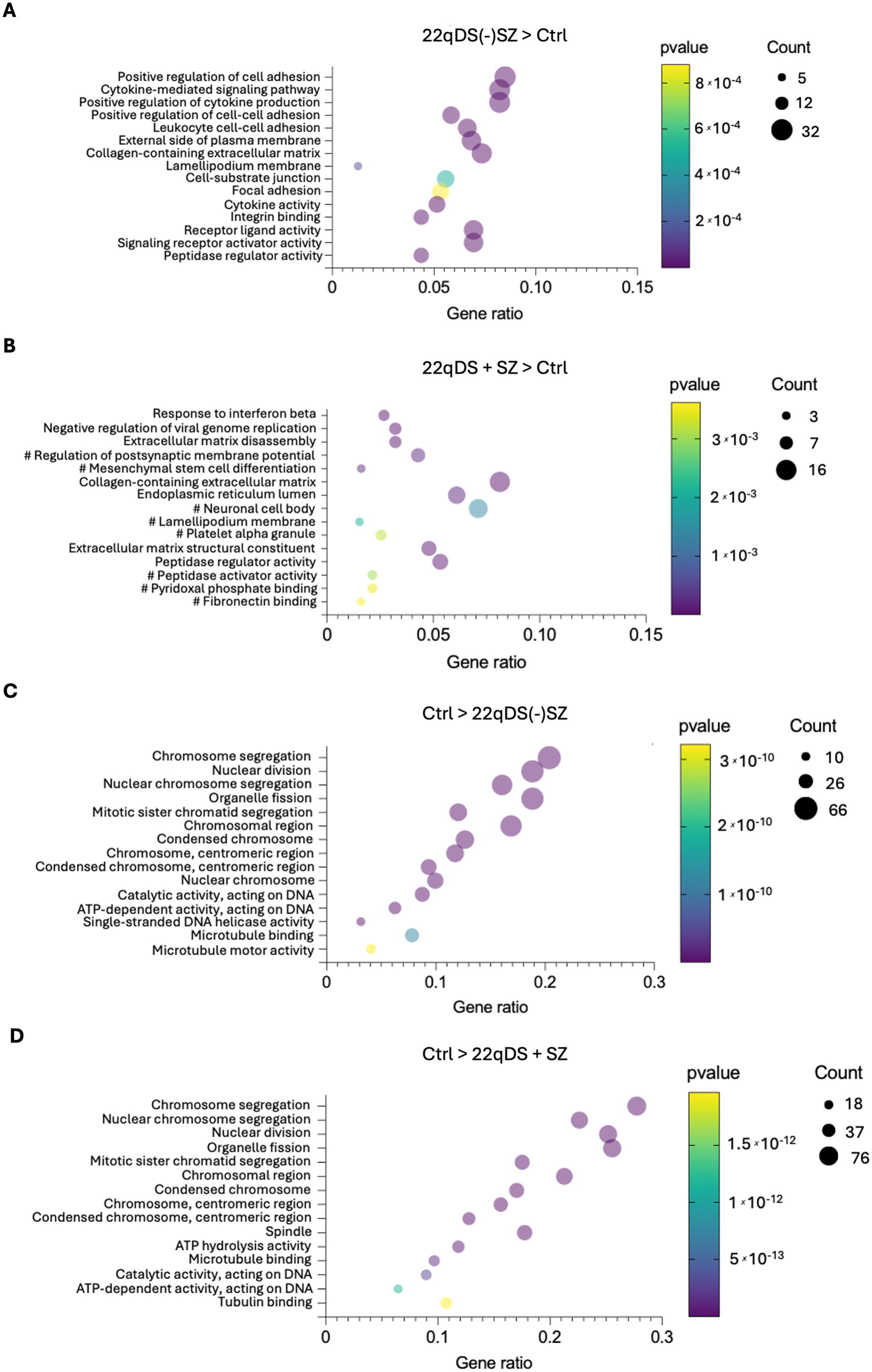
Additional GO term group comparisons. The top 5 significant gene ontology (GO) terms from biological process, cell compartment, and molecular function, respectively (**A**) Enriched in 22qDS(−)SZ over Controls, (**B**) 22qDS + SZ over controls, (**C**) Controls over 22qDS(−)SZ, and (**D**) Controls over 22qDS + SZ. #; GO terms which did not also fall below *p-adjusted value* of 0.1. *n* = 3-5 cell lines per group,

## Notes

### Competing Interest Statement

The authors have declared no competing interest.

